# Metabolic regulation by prostaglandin E_2_ impairs lung group 2 innate lymphoid cell responses

**DOI:** 10.1101/2021.12.23.474031

**Authors:** Calum T. Robb, You Zhou, Jennifer M. Felton, Birong Zhang, Marie Goepp, Privjyot Jheeta, Danielle J. Smyth, Richard M. Breyer, Shuh Narumiya, Henry J. McSorley, Rick M. Maizels, Jürgen K.J. Schwarze, Adriano G. Rossi, Chengcan Yao

## Abstract

Group 2 innate lymphoid cells (ILC2s) play a critical role in asthma pathogenesis. Non-steroidal anti-inflammatory drug (NSAID)-exacerbated respiratory disease (NERD) is associated with reduced signaling via EP2, a receptor for prostaglandin E_2_ (PGE_2_). However, the respective roles for the PGE_2_ receptors EP2 and EP4 (both share same downstream signaling) in the regulation of lung ILC2 responses has yet been deciphered. Here, we find that deficiency of EP2 rather than EP4 augments IL-33-induced lung ILC2 responses and eosinophilic inflammation *in vivo*. In contrast, exogenous agonism of EP4 but not EP2 markedly restricts IL-33- and *Alternaria alternata*-induced lung ILC2 responses and eosinophilic inflammation. Mechanistically, PGE_2_ directly suppresses IL-33-dependent ILC2 activation through the EP2/EP4-cAMP pathway, which downregulates STAT5 and MYC pathway gene expression and ILC2 energy metabolism. Blocking glycolysis diminishes IL-33-dependent ILC2 responses in mice lacking endogenous PG synthesis but not in PG-competent mice. Together, we have defined a mechanism for optimal suppression of lung ILC2 responses by endogenous PGE_2_-EP2 signaling which underpins the clinical findings of defective EP2 signaling in patients with NERD. Our findings also indicate that exogenously targeting the PGE_2_-EP4-cAMP and energy metabolic pathways may provide novel opportunities for treating ILC2-initiated lung inflammation in asthma and NERD.

**Graphical abstract:** 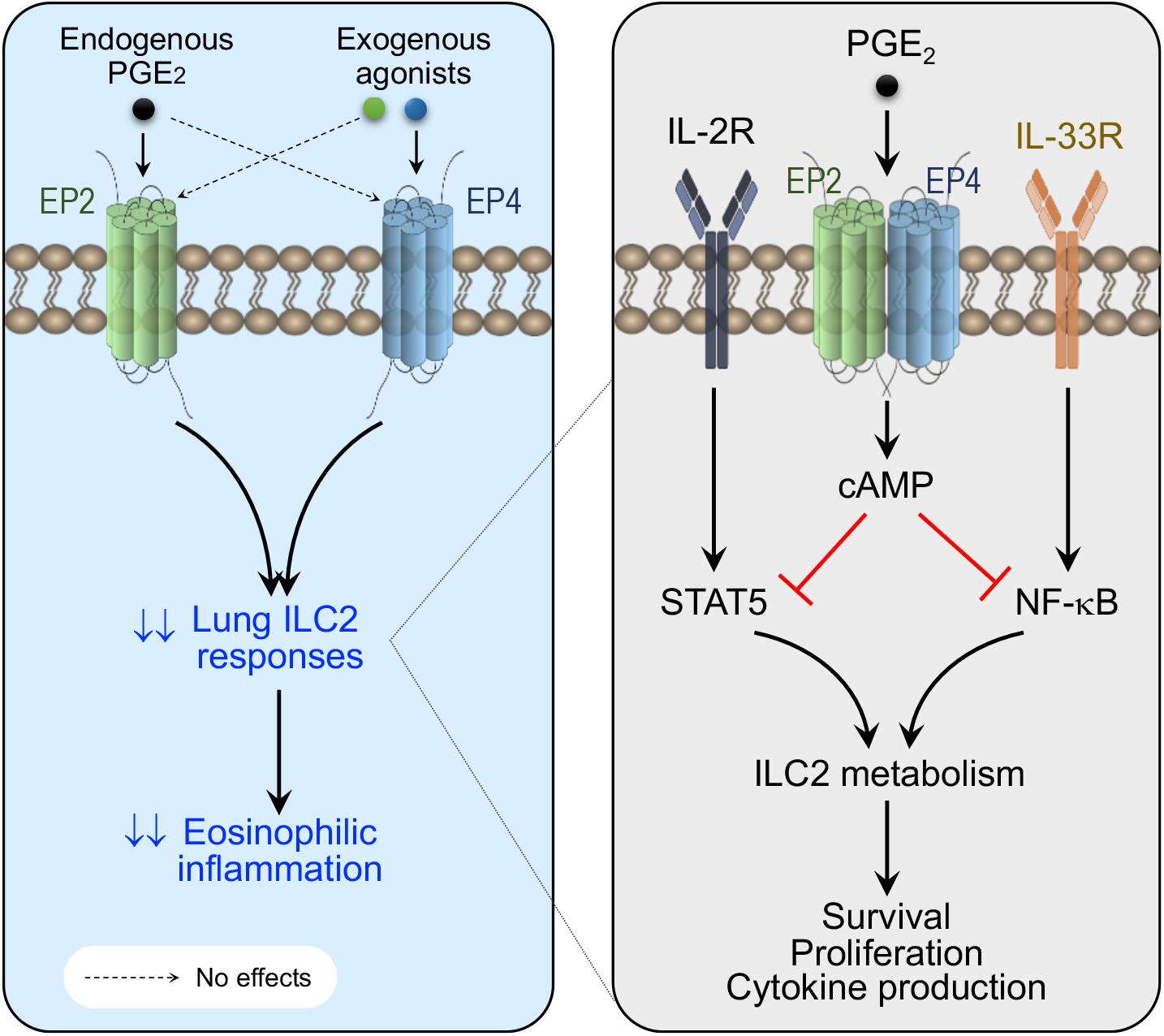

Schematic of potential roles for activation of EP2 and EP4 by endogenous versus exogenous ligands in regulation of lung ILC2 immune responses. Endogenous PGE_2_ in the lung preferentially activates EP2 rather than EP4 to inhibit ILC2 responses and eosinophilic inflammation, and ablation of EP2 enhances lung ILC2 responses. Conversely, lung ILC2 responses are not altered by EP4 deficiency. However, they are markedly inhibited by EP4 agonism but not EP2 agonism. Mechanistically, PGE_2_-EP2/EP4 signaling activates the cAMP pathway which inhibits ILC2 energy metabolism, possibly through interruption of NF-κB (reported in Nagashima H, et al. *Immunity* 2019;51:682-695) and STAT5 signaling, leading to decline of ILC2 survival, proliferation and type 2 cytokine production.

## Introduction

Asthma is a chronic inflammatory lung disease characterized by bronchoconstriction and airway hyperresponsiveness. Upon contact with allergens, irritants (e.g., pollen, air pollutants) or infections, damaged lung epithelial cells release pro-allergic cytokines such as IL-33, which rapidly activates many immune cells including group 2 innate lymphoid cells (ILC2s). ILC2s are innate lymphocytes without antigen-specific receptors, but they highly express Th2 transcription factors such as GATA3 and epithelial cytokine receptors including ST2 (the IL-33 receptor) (1). In response to stimuli by epithelial cytokines, ILC2s produce large amounts of type 2 cytokines (e.g., interleukin [IL]-5, IL-13), which initiate the early onset of innate allergic inflammation (1). Lack or reduction of ILC2s leads to decline of type 2 inflammation during diseases, as found in asthma, metabolic diseases and cancer and alteration of type 2 immunity in response to parasite infections (1)(2). ILC2s also contribute to type 2 helper (Th2) cell activation, sustaining chronic allergic inflammation (3)(4). ILC2s are increased in asthma patients and their cytokine production is associated with disease severity (5)(6).

Non-steroidal anti-inflammatory drugs (NSAIDs) such as aspirin, ibuprofen and indomethacin are widely used for anti-pyretic and analgesic purposes during acute and chronic inflammation through blocking biosynthesis of prostaglandins (PGs) including PGE_2_ (7). However, hypersensitivity reactions to NSAIDs can occur, causing NSAID-exacerbated respiratory disease (NERD), a chronic type 2 immune mediated respiratory disease linked to asthma and nasal polyposis (8). Increased ILC2 numbers have been observed in the nasal mucosa of patients with NERD (9), indicating a role for ILC2s in NERD pathogenesis (10). NERD patients exhibit imbalanced arachidonic acid metabolism with overproduction of PGD_2_ and leukotrienes (e.g., LTD_4_ and LTB_4_) but reduction of PGE_2_ (11). PGD_2_ and leukotrienes promote ILC2 recruitment to the lung and cytokine production (12)(13). Prostacyclin (also called PGI_2_) has been reported to restrict ILC2 activation and effector function (14). NERD patients have reduced expression of the *PTGES* gene (encoding the key PGE_2_ synthase, mPGES1) due to hypermethylation (15). Genetic studies have also suggested that polymorphisms in the *PTGER2* gene (which encodes PGE_2_ receptor EP2) are specifically associated with aspirin-intolerant asthma (16). Moreover, immune cells from NERD patients display reduced EP2 expression (11)(17). These clinical studies suggest that reduction of PGE_2_-EP2 signaling may be a causative factor in the development of NERD, but the underlying mechanisms remain to be elucidated.

PGE_2_ plays distinct roles during the sensitisation and challenge stages of T cell-mediated allergic inflammation, possibly via both EP2 and EP4 (18). PGE_2_ through EP4 suppresses neutrophilic lung inflammation in *ex vivo* human cell culture systems and various animal models (19)(20)(21). Lung allergic responses were increased in mice deficient in PGE_2_ synthases such as COX2 (22). Recently, it was reported that PGE_2_ suppressed human ILC2 function *in vitro* through its receptors EP2 and EP4 (23). However, the respective effects of EP2 and EP4 on ILC2-mediated type 2 immune responses *in vivo* remain unclear, even though EP4 deficiency in haematopoietic cells was reported to increase ILC2 responses in mouse (24). Here, we investigate the impact of endogenous versus exogenous activation of EP2 and EP4 on modulation of ILC2-mediated type 2 immune responses in the mouse lung.

## Results

### Inhibition of endogenous PG synthesis augments IL-33-dependent lung ILC2 responses

At the steady state, PGE_2_ is expressed in most tissues including the lung and its levels are elevated by inflammatory stimuli. We thus investigated whether endogenous PGE_2_ suppressed lung ILC2 immune responses. We administered intratracheally IL-33 into the lungs of wild-type (WT) C57BL/6 mice. Mice also received indomethacin, an NSAID that blocks biosynthesis of all PGs including PGE_2_, or vehicle control in drinking water. In agreement with a previous report (25), indomethacin significantly increased IL-33-mediated accumulation of CD45^+^Lineage^−^ST2^+^ ILC2s to the lung and enhanced type 2 cytokine (i.e., IL-5 and IL-13) production from ILC2s (**Fig 1A,B**). However, indomethacin treatment had no effect on the accumulation of eosinophils in the lung (data not shown), possibly because indomethacin inhibits all PG synthesis and some PGs have direct actions on eosinophils. These results suggest that the net effects of all endogenous PGs suppress lung ILC2 responses. Besides working as a pan-COX inhibitor, indomethacin is also reported to bind to mouse CRTH2 (26), a receptor for prostaglandin D_2_ that mediates ILC2 migration (27)(12). It has been suggested that CRTH2 mediates ILC2 recruitment to the lung when IL-33 is administered systemically (e.g., via intraperitoneal injection), but it has few effects on ILC2 migration to the lung if IL-33 is administered directly to the lung (13). In our studies, we intratracheally administered IL-33 into the lung, thus the enhancement of lung ILC2 responses by indomethacin is unlikely due to its activation of CRTH2, but possibly through inhibition of endogenous PGs.

**Figure 1.**
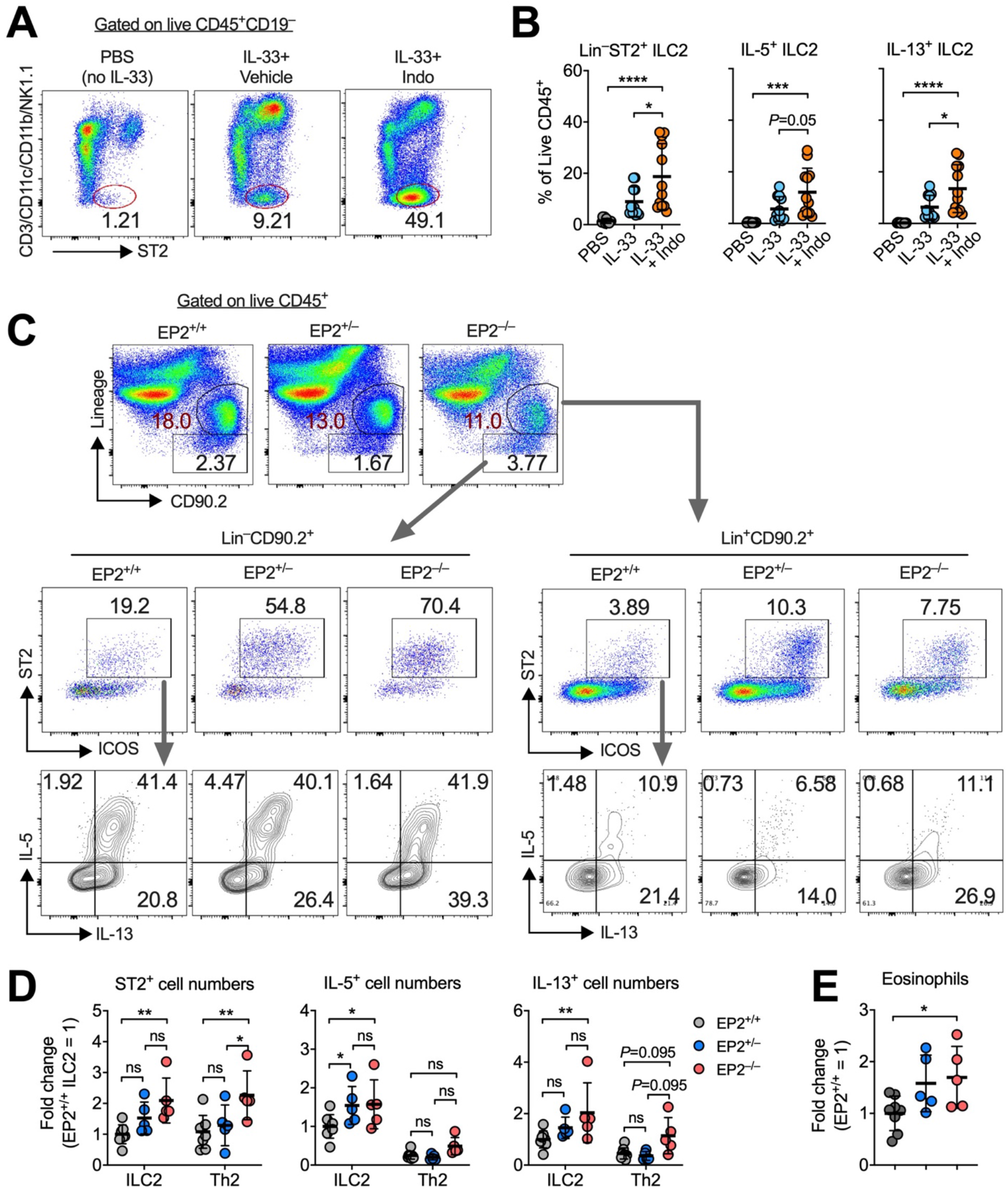
EP2 deficiency augments IL-33-induced lung ILC2 responses. **(A and B)** Representative flow cytometric dot plots **(A)** and collective percentages **(B)** of lung ILC2s in C57BL/6 mice administered intratracheally with PBS (n=11), IL-33 with vehicle (n=11) or indomethacin (n=12) for 3 consecutive days. Cells in **(A)** were pre-gated on live CD45^+^CD19^−^ immune cells. **(C)** Representative flow cytometric dot plots of lineage(CD3/CD19/CD11b/CD11c/NK1.1)^−^CD19^−^ CD90.2^+^ST2^−^ ILC2s and lineage^+^CD90.2^+^ST2^+^ Th2 cells and IL-5 versus IL-13 expression in the lungs of EP2^+/+^ (n=8), EP2^+/–^ (n=5) or EP2^−/–^ (n=5) mice that were administered intratracheally with IL-33 for 3 consecutive days. Cells were pre-gated on live CD45^+^ immune cells. **(D and E)** Number of ST2^+^ total ILC2s, IL-5^+^ ILC2s, IL-13^+^ ILC2s and eosinophils in the lung. Data were normalised by sexes and presented as fold change to the EP2^+/+^ group. Data shown as means ± SDs were pooled from three independent experiments. Each dot in the bar graphs represents one mouse. \**P*<0.05, \*\**P*<0.01, \*\*\**P*<0.001, \*\*\*\**P*<0.0001 by one-way ANOVA with post-hoc Holm-Sidak’s multiple comparisons tests. ns, not significant.

### EP2 deficiency enhanced on IL-33-induced lung ILC2 responses

As indomethacin inhibits biosynthesis of all PGs, we examined whether PGE_2_ is involved in regulation of lung ILC2 responses, and if yes, through which receptor? We examined published datasets (28) and found that ILC2s isolated from various tissues including lung, bone marrow, skin and intestine have considerably higher gene expression of EP4, followed by EP2 and EP1, and that EP3 has much lower expression levels in ILC2s (**Fig S1**). We first asked whether blockade of endogenous PGE_2_ signaling via the EP2 receptor influenced lung ILC2 responses. We injected IL-33 into the lungs of EP2-deficient (29) or control mice and measured lung ILC2 activation. As expected, EP2-deficiency increased lung ILC2 accumulation and production of type 2 cytokines (IL-5 and IL-13) in response to IL-33 (**Fig 1C,D**). EP2 deficiency also increased lung Lin^+^CD90.2^+^ST2^+^ T cells, but type 2 cytokine production from T cells was not significantly affected by loss of EP2 (**Fig 1C,D**). In agreement with the increase in ILC2s, EP2 deficient mice had more eosinophils after IL-33 treatment (**Fig 1E**). Our data suggest that endogenous PGE_2_-EP2 signaling restricts IL-33-dependent lung ILC2 responses.

### EP4 deficiency does not affect IL-33-induced lung ILC2 responses

To test whether deleting EP4 signaling modulates lung ILC2 responses, we administered IL-33 to global EP4-deficient (30) and control mice. Unexpectedly, EP4 deficiency did not enhance ILC2 accumulation in the lung but reduced type 2 cytokine expression from lung ILC2s compared to WT and heterozygous control mice (**Fig 2A,B**). Global EP4 deficiency also did not increase eosinophils in the lung (**Fig 2C**). Thus, we then investigated whether specific deletion of EP4 in ILC2s enhances lung ILC2 responses.

**Figure 2.**
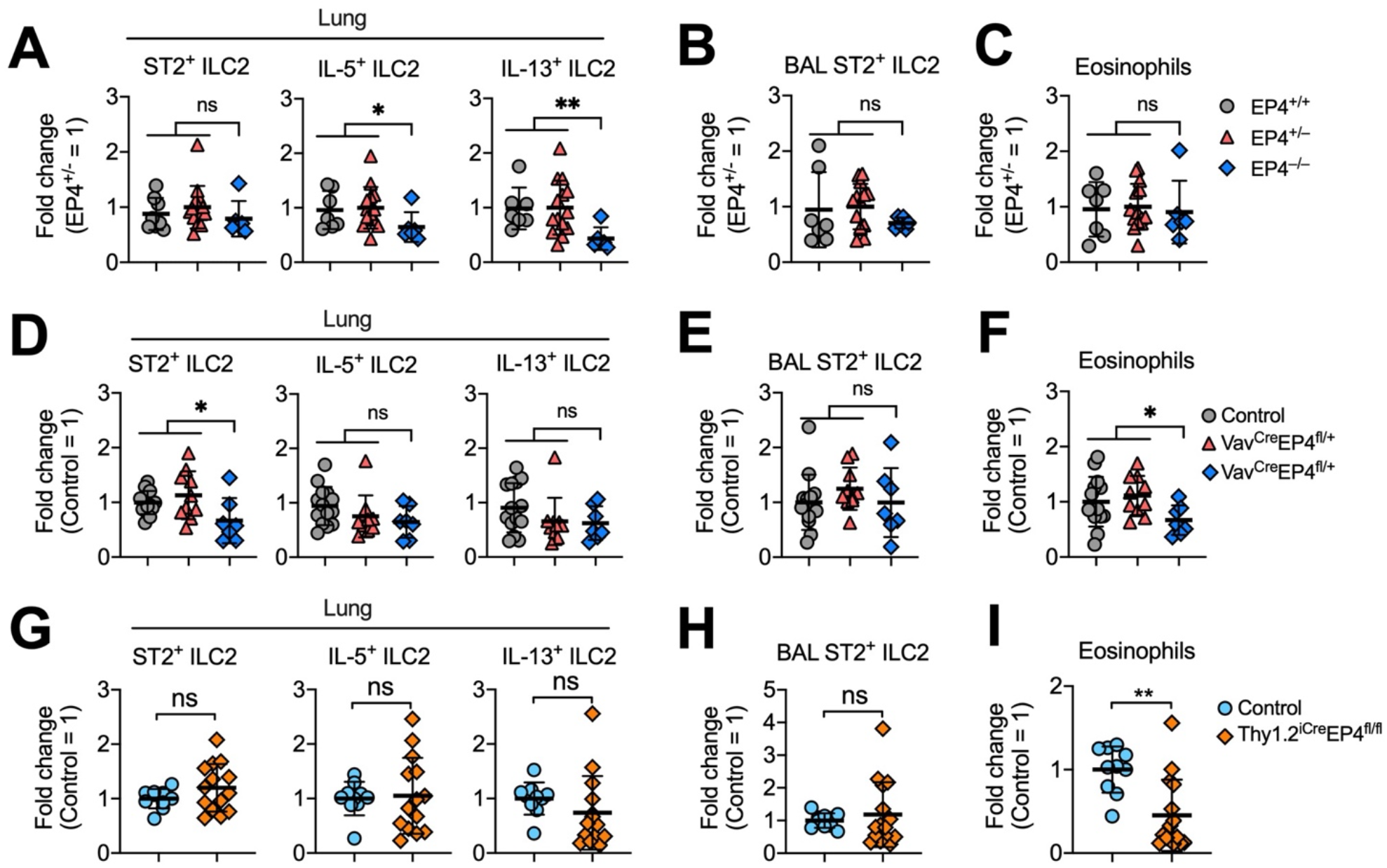
Endogenous EP4 signaling is dispensable for the control of IL-33-induced lung ILC2 responses. **(A to C)** Number of lung ST2^+^ total ILC2s, IL-5^+^ ILC2s and IL-13^+^ ILC2s (A), BAL ST2^+^ ILC2s **(B)** and eosinophils **(C)** in EP4^+/+^ (n=7), EP4^+/–^ (n=14) or EP4^−/–^ (n=6) mice that were administered intratracheally with IL-33 for 3 consecutive days. **(D to F)** Number of lung ST2^+^ total ILC2s, IL-5^+^ ILC2s and IL-13^+^ ILC2s **(D)**, BAL ST2^+^ ILC2s **(E)** and eosinophils **(F)** in control (EP4^fl/fl^ or EP4^fl/+^, n=14), Vav^Cre^EP4^fl/+^ (n=10), or Vav^Cre^EP4^fl/fl^ (n=7) mice that were administered intratracheally with IL-33 for 3 consecutive days. **(G to I)** Number of lung ST2^+^ total ILC2s, IL-5^+^ ILC2s and IL-13^+^ ILC2s **(G)**, BAL ST2^+^ ILC2s **(H)** and eosinophils **(I)** in control (EP4^fl/fl^ or EP4^fl/+^, n=10) or Thy1.2^iCre^EP4^fl/fl^ (n=14) mice that were administered intratracheally with IL-33 for 3 consecutive days. Data were normalised by sexes and presented as fold changes to the EP4^+/–^ **(A to C)** or control **(D to I)** groups, respectively. Data shown as means ± SDs were pooled from three independent experiments. Each dot in the bar graphs represents one mouse. \**P*<0.05, \*\**P*<0.01 by one-way ANOVA with *post-hoc* Holm-Sidak’s multiple comparisons tests **(A to F)** or unpaired, 2-tailed Student’s t-test **(G to I)**. ns, not significant.

As there was no mouse lines that selectively targets only ILC2s, we generated Vav^Cre^EP4^fl/fl^ mice by crossing EP4-floxed mice (31) to Vav-cre mice (32) which drives ablation of EP4 in all hematopoietic-derived immune cells (including ILC2s). A previously published report showed that Vav-dependent EP4 deficiency enhanced lung ILC2 function and eosinophilic inflammation (24). However, we found that conditional EP4 deficient mice also had comparable lung ILC2 responses in response to IL-33 administration, albeit with a moderate reduction of lung ST2^+^ ILC2 accumulation (**Fig 2D,E**). Homozygous Vav^Cre^EP4^fl/fl^ mice had reduced eosinophils (**Fig 2F**), which was likely due to a direct effect of EP4 on Vav-expressing eosinophils. Both EP4^−/–^ and Vav^Cre^EP4^fl/fl^ mice ablated EP4 from the germline, which may have a potential effect on ILC (including ILC2) development. To exclude this possibility, we employed another inducible EP4 deficient mouse line by crossing Thy1.2^iCre^ mice(33) to EP4-floxed mice to generate Thy1.2^iCre^EP4^fl/fl^ mice, which don’t have EP4 expression in Thy1.2-expressing cells (i.e., all T cells and ILCs) after administration of tamoxifen. Again, Thy1.2-Cre driven inducible EP4 deficiency did not alter IL-33-induced lung ILC2 responses although it significantly reduced lung eosinophils (**Fig 2G-I**). These results suggest that endogenous PGE_2_-EP4 signaling does not significantly affect lung ILC2 responses.

### EP4 agonism inhibits the alarmin IL-33- and fungal antigen-induced lung ILC2 responses

To examine the effects of exogenous activation of EP2 and EP4 on ILC2 responses *in vivo*, we intratracheally administered IL-33 into C57BL/6 mice. Administration of IL-33 induced accumulation of eosinophils and neutrophils in the bronchoalveolar lavage (BAL) and lung, whilst such accumulation was almost completely attenuated by co-administration of an EP4 agonist (**Fig 3A-C**). Co-administration of an EP2 agonist had no significant effects on IL-33-dependent accumulation of eosinophils and neutrophils (**Fig 3A-C**). Flow cytometric analysis suggested that IL-33-induced recruitment of lineage-negative ST2^+^ ILC2s in the airway was also impeded by the EP4 agonist rather than the EP2 agonist (**Fig 3D,E**). Analysis of lung single cells showed that IL-33 increased numbers of total ST2^+^ ILC2s and their production of type 2 cytokines (e.g., IL-5 and IL-13), with accumulation of ILC2s was again inhibited by the EP4 (but not EP2) agonist (**Fig 3F-I**).

**Figure 3.**
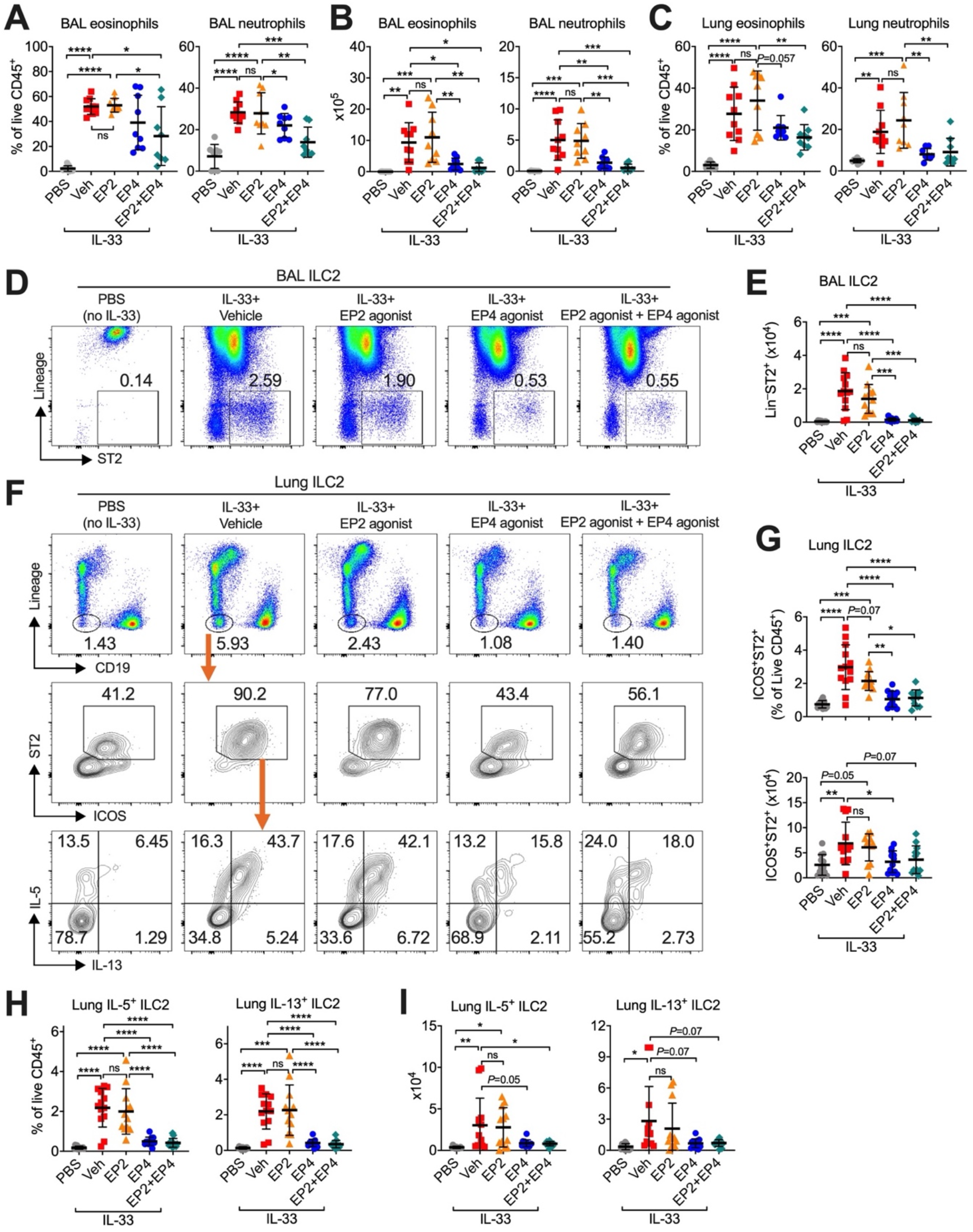
EP4 agonism inhibits the alarmin IL-33-activated lung ILC2 responses. C57BL/6 mice were administered intratracheally with PBS (n=12) or IL-33 with vehicle (n=13), EP2 agonist (butaprost, n=12), EP4 agonist (L-902,688, n=12), or both EP2 and EP4 agonists (n=11) for 3 consecutive days. **(A to C)** Eosinophils and neutrophils in the lungs and the bronchoalveolar lavages (BAL) fluids. **(D to E)** Representative flow cytometric dot plots and collective numbers of lineage(CD3/CD19/CD11c/CD11b/NK1.1)^−^ST2^+^ ILC2s in the BAL fluid. Cells in **(D)** were pre-gated on the live CD45^+^ immune cells. **(F)** Representative flow cytometric dot plots of lineage(CD3/CD11c/CD11b/NK1.1)^−^CD19^−^, ST2^+^ ILC2s, and IL-5-versus IL-13-expressing ILC2s in the lungs. Cells were pre-gated on the live CD45^+^ immune cells. **(G to I**) Collective percentages and numbers of ST2^+^ total ILC2s and IL-5- or IL-13-expressing ILC2s in the lungs. Data shown as means ± SDs were pooled from 2-3 independent experiments. Each dot in the bar graphs represents one mouse. \**P*<0.05, \*\**P*<0.01, \*\*\**P*<0.001, \*\*\*\**P*<0.0001 by one-way ANOVA with *post-hoc* Holm-Sidak’s multiple comparisons tests. ns, not significant.

Next, we used an IL-33-dependent acute lung inflammation model induced by extracts of *Alternaria alternata* (A.A.), an airborne fungus that can promote the development of allergic diseases like asthma and eczema (34). Similarly, co-administration of the EP4, but not EP2, agonist significantly diminished IL-13 production from lung ILC2s (**Fig 4A-C**). Of note, EP4 agonist alone did not reduce IL-5 expression in ILC2s, possibly due to the short-term treatment (24 h) (**Fig 4A-C**). However, co-administration of EP2 and EP4 agonists reduced both IL-5 and IL-13 production from ILC2s at the single cell level (**Fig 4D**). Consistently, exogenous activation of EP4 or both EP2 and EP4 markedly prevented A.A.-dependent accumulation of eosinophils to the lung (**Fig 4E**). These results indicate that exogenous activation of EP4 effectively limits ILC2 responses and lung inflammation.

**Figure 4.**
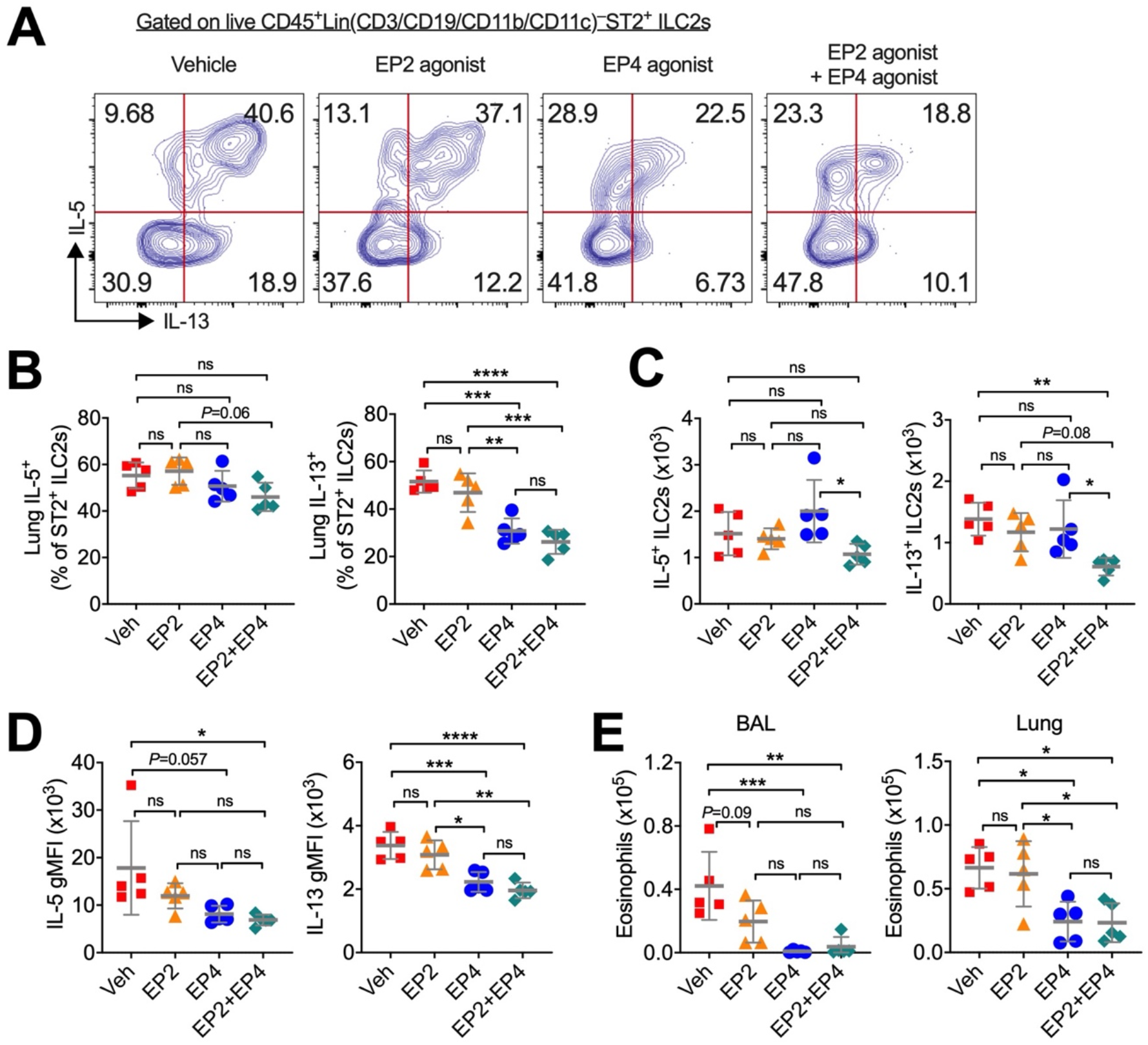
Exogenous activation of EP4 inhibits *Alternaria alternata* induced lung ILC2 responses. C57BL/6 mice were administered intratracheally with *Alternaria alternata* allergen (A.A.) with vehicle (n=5), EP2 agonist (butaprost, n=5), EP4 agonist (L-902,688, n=5), or both EP2 and EP4 agonists (n=5) for 24 h. **(A)** Representative flow cytometric dot plots of IL-5-versus IL-13-expressing ILC2s in the lungs. Cells were pre-gated on the live CD45^+^lineage(CD3/CD19/CD11b/CD11c/NK1.1)^−^ST2^+^ ILC2s. **(B and C)** Collective percentages and numbers of IL-5^+^ or IL-13^+^ ILC2s in the lungs. **(D)** Geometric median fluorescence intensity (gMFI) of IL-5 and IL-13. **(E)** Numbers of eosinophils in the lungs. Data were shown as means ± SDs. Each dot in the bar graphs represents one mouse. \**P*<0.05, \*\**P*<0.01, \*\*\**P*<0.001, \*\*\*\**P*<0.0001 by one-way ANOVA with post-hoc Holm-Sidak’s multiple comparisons tests. ns, not significant.

### PGE_2_-EP2/EP4-cAMP signaling directly inhibits ILC2 activation *in vitro*

We next examined the underlying mechanisms for PGE_2_ suppression of ILC2 responses *in vitro*. To this end, we stimulated whole lung or bone marrow cells isolated from Rag2^−/−^ mice (which have no T or B cells) *in vitro* with IL-33, IL-2 and IL-7. This cytokine cocktail induced production of type 2 cytokines in the supernatants after 3-5 days of culture (**Fig 5A,B**). As seen in human ILC2 cell cultures (23), addition of exogenous PGE_2_ significantly reduced IL-33-induced IL-5 and IL-13 production (**Fig 5A,B**). Blocking the synthesis of endogenous PGs with indomethacin had no effects on IL-5 and IL-13 production from lung cell cultures (**Fig 5A**) but significantly upregulated IL-13 production from whole bone marrow cell cultures (**Fig 5B**). The inhibitory effect of PGE_2_ on type 2 cytokine production was mimicked by an EP2 agonist and an EP4 agonist (**Fig 5A**). Results from flow cytometric analysis also showed that PGE_2_, EP2 agonist and EP4 agonist suppressed survival and expansion of total ST2^+^ ILC2s and their expression of IL-5 and IL-13 (**Fig S2**). Furthermore, the inhibitory effect of PGE_2_ was also mimicked by dibutyryl-cyclic AMP (db-cAMP, a cell-permeable cAMP analog) and IBMX (an inhibitor of cyclic nucleotide phosphodiesterases that increases the intracellular cAMP level through blocking cAMP degradation) (**Fig 5B**). These results strongly suggest that the PGE_2_-EP2/EP4-cAMP signaling pathway suppresses ILC2 responses *in vitro*.

**Figure 5.**
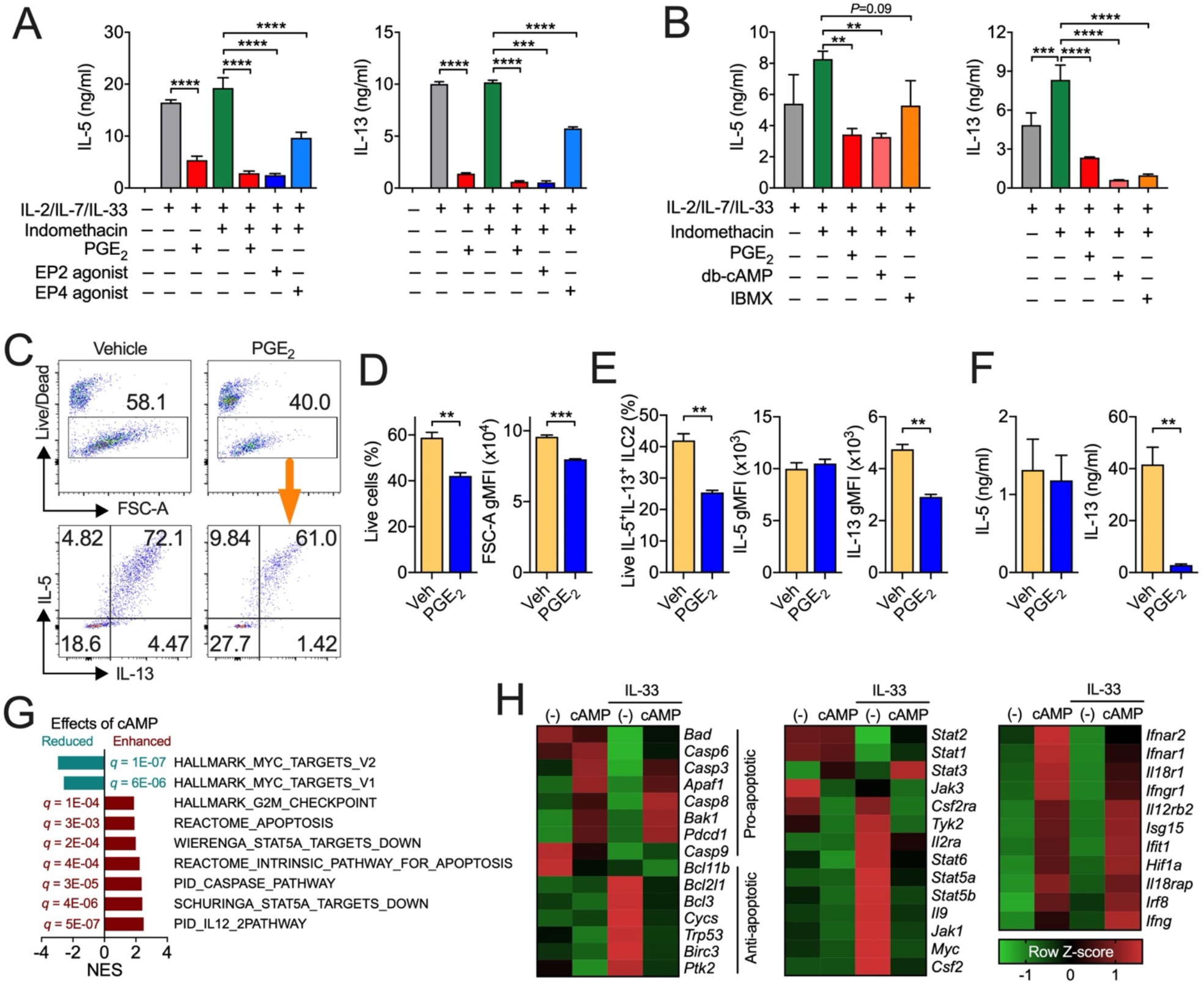
PGE_2_-cAMP signaling directly inhibits ILC2 activation and alters ILC2 gene expression associated with cellular metabolic pathways. (**A and B)** IL-5 and IL-13 production from whole lung **(A)** or bone marrow **(B)** cells from Rag2^−/–^ mice that were cultured *in vitro* with IL-2, IL-7 and IL-33 in the presence or absence of indicated reagents. Data shown as means ± SEMs are from one of two independent experiments. \**P*<0.05, \*\**P*<0.01, \*\*\**P*<0.001, \*\*\*\**P*<0.0001 by one-way ANOVA with post-hoc Holm-Sidak’s multiple comparisons tests. **(C to F)** Representative flow cytometric dot plots **(C)**, cell viability and size (FSC-A) **(D)**, collective percentages of live IL-5^+^IL-13^+^ ILC2s and geometric fluorescence intensity (gMFI) of IL-5 and IL-13 (F), and cytokine secretion in the supernatants **(F)** of ILC2s sorted from Rag2^−/–^ lungs and then cultured *in vitro* with IL-33, IL-2 and IL-7 for 3 days in the presence or absence of PGE_2_. Data shown as means ± SEMs are from one of two independent experiments. \*\**P*<0.01, \*\*\**P*<0.001 by unpaired, 2-tailed Student’s *t*-test. Veh, vehicle; FSC-A, forward scatter area. **(G to H)** Gene enrichment assays **(G)** and changes of gene expression of pathways of interest **(H)** in ILC2s cultured with IL-33 and/or cAMP for 4 h. Raw RNAseq data were retrieved from Gene Expression Omnibus GSE131996, reanalysed and transformed to Z-scores. NES, normalized enrichment score.

To investigate whether PGE_2_ directly controls ILC2 activation, we sorted ILC2s from lung (**Fig 5C-F**) or bone marrows (**Fig S3**) and cultured them with IL-33 and/or PGE_2_. Addition of PGE_2_ significantly decreased ILC2 viability, cellular size (i.e., FSC [forward scatter]), and IL-13 (but not IL-5) expression and secretion (**Fig 5C-F and Fig S3**). This is in agreement with a previous report showing that cAMP inhibits ILC2 production of IL-13 rather than IL-5 (35). These results suggest that PGE_2_ directly acts on ILC2s and controls ILC2 survival, growth and function.

### cAMP regulates IL-33-dependent and IL-33-independent ILC2 gene expression

To understand the underlying mechanisms for PGE_2_ suppression of ILC2 responses, we examined a published dataset (35) and analysed the effects of cAMP on ILC2 gene expression. Gene set enrichment analysis showed that, in addition to restricting cell cycle progression as reported previously (35), cAMP markedly enhanced apoptosis-associated pathways (**Fig 5G**). Expression of pro-apoptosis associated genes (e.g., *Bad, Bak1*, caspases, *Pdcd1*) were upregulated by cAMP independently of IL-33 (**Fig 5H**). In contrast, IL-33-induced expression of anti-apoptotic genes (e.g., *Bcl2l1, Bcl3, Trp53, Ptk2*) were down-regulated by cAMP (**Fig 5H**). This differential cAMP-altered expression of pro- and anti-apoptotic genes contributes to overall reduced cell survival. Moreover, IL-33 elevated expression of the STAT5 pathway genes including cytokines/receptors (*Il2ra, Il9, Csf2, Csf2r*), JAK family members (*Jak1, Tyk2*) and *Stat5a* and *Stat5b* themselves, which were all down-regulated by cAMP (**Fig 5H**). STAT5 is important for ILC2 accumulation at lymphoid and nonlymphoid tissues (36). Similar to findings observed in T cells (37), cAMP up-regulated gene expression involved in the IL-12 pathway (e.g., *Il12rb2*) and the interferon pathway, the latter included cytokine receptors (*Il18r, Il18rap, Ifng, Ifngr1, Ifnar1, Ifnar2*) and downstream interferon-stimulated or gamma-activated genes (*Ifit1, Irf8, Isg15, Hif1a*) (**Fig 5H**). IL-33 had no impacts on IL-12 and interferon pathway gene expression (**Fig 5H**). The interferon- and IL-12-activated STAT1/2/4 repress ILC2 activation by antagonizing STAT5 (38). Similarly, cAMP enhanced STAT3 expression (**Fig 5H**) which usually competes with STAT5 for binding to gene promoters (39). Furthermore, STAT5 is recruited to the c-Myc enhancer (40), which may contribute to cAMP down-regulation of the MYC pathway (**Fig 5G**). MYC plays an important role in cellular metabolism, and it has been reported to play a critical role in ILC2-mediated airway inflammation (41).

### PGE_2_ regulates ILC2 cellular metabolism

In this study, we have established that PGE_2_ reduces ILC2 survival, cell size and proliferation **(Fig 5C,D**), all of which are hallmark events during cellular metabolism. Similarly, *in vivo* activation of EP4 and, possibly EP2, by their respective agonists reduced Ki-67-expressing ST2^+^ ILC2s in lungs of mice treated with IL-33 (**Fig 6A,B**). In contrast, EP2 deficiency increased Ki-67^+^ST2^+^ ILC2s in the lung (**Fig 6C,D**). These results, together with the results from RNA-seq analysis (**Fig 5G,H**), may imply changes in energy metabolism of ILC2s by PGE_2_-cAMP signaling. Therefore, to examine if PGE_2_ directly regulates ILC2 metabolism, we measured metabolic changes in cultured lung ILC2s using the Seahorse assays. Indeed, PGE_2_ reduced oxygen consumption rate (OCR, an indicator of mitochondrial phosphorylation), extracellular acidification rate (ECAR, an indicator of aerobic glycolysis) and glycolytic proton efflux rate (PER, a real-time indicator of glycolysis rate) of ILC2s (**Fig 6E-G**), suggesting that PGE_2_ directly inhibits IL-33-dependent ILC2 energy metabolism *in vitro*.

**Figure 6.**
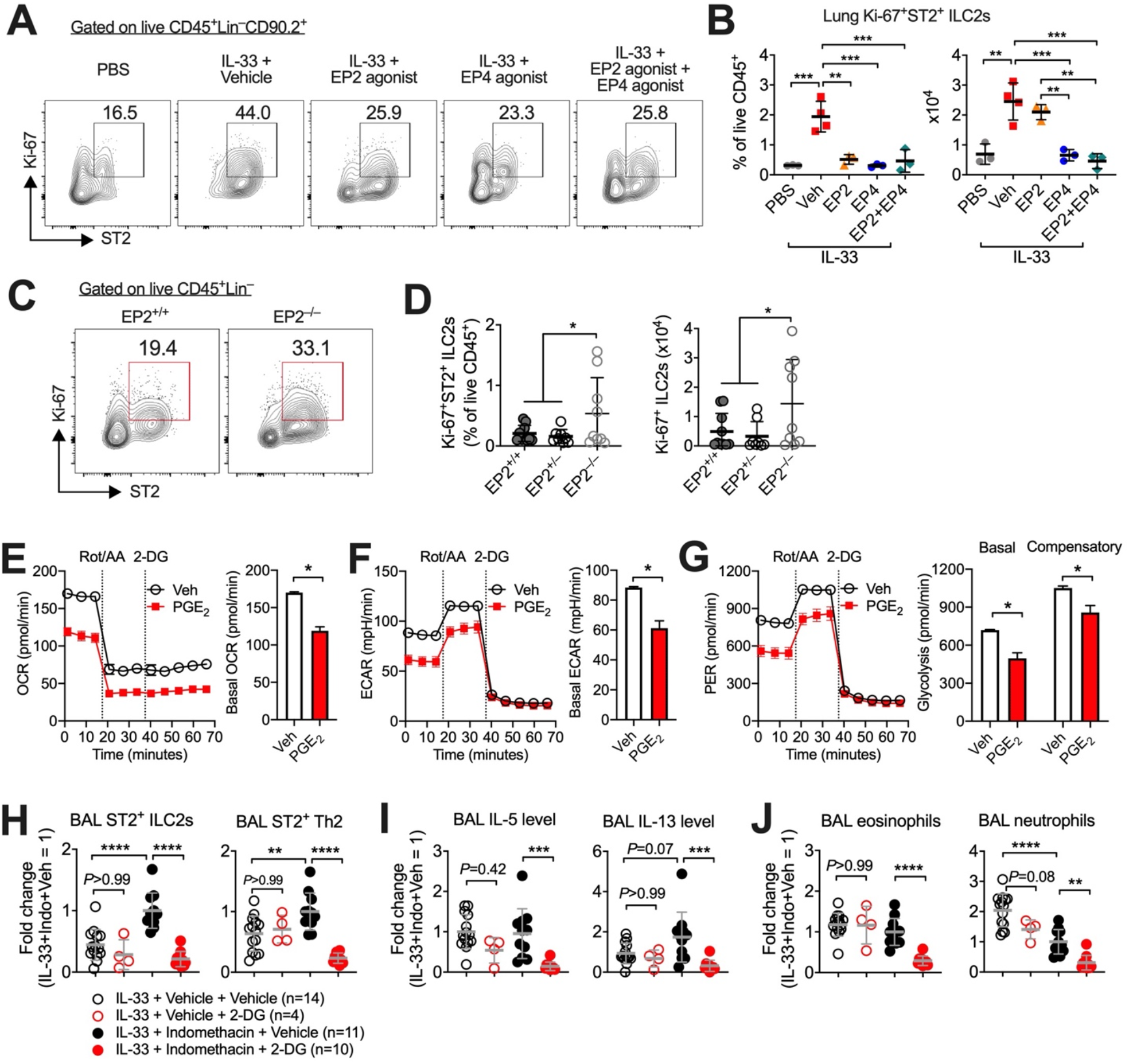
PGE_2_ represses ILC2 cellular metabolism. **(A and B)** Representative flow cytometric dot plots **(A)** and collective percentages and numbers **(B)** of lung proliferating Ki-67^+^ ILC2s in C57BL/6 mice administered intratracheally with IL-33 with PBS (n=3) or IL-33 with vehicle (n=4), an EP2 agonist (butaprost, n=3), an EP4 agonist (L-902,688, n=3), or both the EP2 and EP4 agonists (n=3) for 3 consecutive days. **(C and D)** Representative flow cytometric dot plots **(C)** and collective percentages and numbers **(D)** of lung proliferating Ki-67^+^ ILC2s in EP2^+/+^ (n=10), EP2^+/–^ (n=8) or EP2^−/–^ (n=9) mice administered with IL-33 for 3 consecutive days. **(E to G)** The oxygen consumption rate (OCR), extracellular acidification rate (ECAR), and proton efflux rate (glycoPER) of ILC2s that were sorted from lungs of Rag2^−/–^ mice and then cultured with IL-33 with or without PGE_2_ for 3 days. **(H to J)** Numbers of ST2^+^ ILC2 and ST2^+^ Th2 cells **(H)**, IL-5 and IL-13 levels **(I)**, and eosinophil and neutrophil numbers **(J)** in bronchoalveolar lavages (BAL) mice administered intratracheally with IL-33 with or without 2-DG for 3 consecutive days. Numbers of mice in each group are indicated in the keys. Indomethacin or its vehicle (DMSO) was administered through drinking water. Data shown as means ± SDs **(B, D, H to J)** or means ± SEMs **(C to E)** were pooled from one **(B)**, two **(D)** or three **(H to J)** experiments or representative one of two experiments **(E to G)**. Data in **(H to J)** were normalized by sexes and presented as fold changes to the IL-33+Indomethacin+vehicle group. Each dot in the bar graphs **(B, D, H to J)** represents one mouse. \**P*<0.05, \*\**P*<0.01, \*\*\**P*<0.001, \*\*\*\**P*<0.0001 by one-way **(B)** or two-way **(H to J)** ANOVA with post-hoc Holm-Sidak’s multiple comparisons tests or unpaired, 2-tailed Student’s *t*-tests **(D to G)**.

To determine whether PG signaling controls ILC2 responses through inhibiting ILC2 metabolism in vivo, we treated WT C57BL/6 mice with IL-33 and indomethacin. Some mice received 2-deoxy-D-glucose (2-DG), a glucose analogue that inhibits glycolysis through blocking glucose hexokinase. Similar to previous reports showing that 2-DG had no impact on IL-33- or helminth infection-induced ILC2 activation (42)(43), here we also report that 2-DG affected neither IL-33-induced ILC2 accumulation and type 2 cytokine production nor eosinophil recruitment in mice with intact PG signaling (i.e., without indomethacin-treatment) (**Fig 6H-J**). However, in indomethacin-treated mice where biosynthesis of endogenous PGs including PGE_2_ was blocked, 2-DG strikingly reduced ILC2 and Th2 numbers and type 2 cytokine production in the lung (**Fig 6H-I**). Accordingly, overall lung inflammation was also decreased by 2-DG in indomethacin-treated mice as evidenced by reduction of both eosinophils and neutrophils (**Fig 6J**). These results suggest that PGE_2_ impedes ILC2 energy metabolism and constricts IL-33-induced type 2 immune responses.

## Discussion

Here, we report that PGE_2_-EP2/EP4 signaling limits type 2 lung inflammation through negative regulation of ILC2 responses. Both EP2 and EP4 activate the downstream cAMP-PKA signaling pathway, thus can have redundant or additive functions *in vivo*. Blockade of both receptors is more effective than blocking just one receptor. For example, EP2 and EP4 collaboratively promotes T cell-mediated psoriasis (44). Under certain circumstances, the actions of PGE_2_ *in vivo* may be dominantly mediated by one receptor, leaving the other receptor dispensable. For example, the effects of PGE_2_ on T cell-mediated intestinal homeostasis and gut or brain inflammation are chiefly mediated by EP4, and EP2 exerts few effects (45)(46)(47). In this study, we have found distinct effects of EP2 and EP4 that dependent on whether they are stimulated by endogenous PGE_2_ or by exogenous agonists. While the lung ILC2 responses were enhanced by EP2 deficiency, it was not influenced by exogenous EP2 agonism. In contrast, while ablation of EP4 had little impact on lung ILC2 responses and eosinophilic inflammation, these were strongly suppressed by exogenous activation of EP4. These findings suggest that in the lung EP2 may have already been endogenously activated at the optimal level for repressing ILC2 responses, leading to unresponsiveness to further EP2 agonism. On the other hand, endogenous PGE_2_ does not seem to significantly activate EP4 signaling, thus allowing for significantly additional activation of EP4 by an exogenous agonist, resulting in considerable effects. The discrepancy in effects between EP4 agonism and EP4 deficiency on regulation of lung ILC2 responses may also indicate that EP4 expression on different lung cell types has distinct effects during type 2 inflammation and/or that there are distinct roles for the strength (e.g., the amounts of local lung resident PGE_2_) and the timing for endogenous versus exogenous EP4 activation. Future studies will need to examine these possibilities.

PGE_2_ has been indicated to be associated with various lung conditions including neutrophilic inflammation (19), infections (e.g., mycobacterium tuberculosis (17)) and COPD (21). Our findings that EP2 rather than EP4 is the dominant mediator of endogenous PGE_2_ effects on the regulation of lung ILC2 responses specifically reflect findings from clinical studies. Indeed, *PTGER2* gene polymorphisms are associated with asthma especially NERD and patients with NERD have decreased EP2 expression (16)(11)(17). Our results suggest that the down-regulation or alteration of EP2 signaling in NERD patients may contribute to augmented ILC2 responses and inflammation. Thus, pharmacological activation of EP4 may be a promising therapeutic approach to limit augmented ILC2 responses in asthma patients with low EP2 expression or function.

We also report that PGE_2_ through activation of cAMP suppresses ILC2 energy metabolism, which controls cell survival, proliferation, activation and effector function. Besides PGE_2_, many molecules that can activate the cAMP pathway such as PGI_2_ (14), β2-adrenergic receptor agonist (e.g., norepinephrine) (48), and calca-encoding calcitonin gene-related peptide (35)(49)(50) have all been found to suppress ILC2 responses and indeed allergic lung inflammation. Our findings of regulation of ILC2 gene expression by the PGE_2_-cAMP pathway via IL-33-dependent and IL-33-independent mechanisms suggest that PGE_2_ may have broad impacts on ILC2-mediated type 2 immune responses. It is worth noting that although cAMP represses ILC2 activation, it does not inhibit IL-5 expression by ILC2s, which was found in this current study and in line with the findings of other studies (35). Indeed, cAMP was reported to mediate elevation of IL-5 production from homeostatic ILC2s (51).

Current treatments do not provide a cure for asthma and in the case of NERD patients, such treatments may not be effective. Our work highlights the EP4-cAMP pathway as a potential therapeutic target for asthma and in particular for NERD, where small molecules that harness activation of EP4 and/or elevation of intracellular cAMP levels could be of clinical benefit. Phosphodiesterase 4 inhibitors, which increase the intracellular cAMP level (via preventing cAMP degradation), have been considered for treatment of asthma and other lung diseases (52)(53). Furthermore, we propose that targeting ILC2 energy metabolic pathways (e.g., glycolysis) may be beneficial in the control of NSAID-dependent augmentation of type 2 lung inflammation in patients with NERD.

## Methods

### Mice

*Rag2*^−/−^, EP2(Ptger2)^−/–^ (29), EP4(*Ptger4*)^−/−^ (30) mice were maintained under specific pathogen-free conditions in accredited animal facilities in the University of Edinburgh. Vav^Cre^EP4^fl/fl^ and Thy1.2^iCre^EP4^fl/fl^ mice were generated by crossing Vav-Cre mice (32) and Thy1.2^iCre^ mice (33), respectively, to EP4-floxed mice and maintained under specific pathogen-free conditions in accredited animal facilities in the University of Edinburgh. Wild-type C57BL/6 mice were purchased from Harlan UK or bred in animal facilities within the University of Edinburgh. Sex-matched mice aged >6 weeks were used in experiments. Mice were randomly allocated into different treatment groups and analysed individually. No animal was excluded for analysis except one in Fig2A which was overlooked at the step of *ex vivo* restimulation of lung single cells with PMA, ionomycin and Brefeld A (for detecting cytokine expression).

### Reagents

Antibodies to mouse CD45 (clone 30-F11), CD19 (clone 6D5), NK-1.1 (clone PK136), CD4 (clone RM4-5), CD90.2 (clone 30-H12), CD3e (clone 145-2C11), CD11b (clone M1/70), CD11c (clone N418), ICOS (clone7E.17G9), IL-5 (clone TRFK5), IL-13 (clone eBio13A), Ki-67 (clone 16A8) and Ly6G (clone 1A8) were purchased from eBioscience or Biolegend. Antibodies to mouse ST2(IL33-R) (clone DJ8) and SiglecF (clone E50-2440) were purchased from mdbioproducts and BD respectively. LIVE/DEAD™ Fixable Blue Dead Cell Stain Kit for UV excitation, UltraComp eBeads™ and mouse IL-5/IL-13 ELISA kits were purchased from Thermo Fisher Scientific. Recombinant mouse IL-33 and *Alternaria alternata* (A.A) extract were purchased from Biolegend and Stallergenes Greer, respectively. PGE_2_ and L-902,688 (EP4 agonist) were obtained from Cayman. Indomethacin, phorbol myristate acetate (PMA), ionomycin, brefeldin A, (R)-Butaprost (EP2 agonist), dibutyryl cyclic adenosine monophosphate (db-cAMP) and 3-isobutyl-1-methyxanthine (IBMX) were purchased from Sigma. Complete RMPI consisted of RPMI 1640 (Gibco) supplemented with 10% FBS, 1% Penicillin/Streptomycin, 1% L. glutamine and 50 μM β-mercaptoethanol.

### Airway administration of pro-allergic cytokines and *Alternaria* allergen

A variety of different mouse lines including C57BL/6 wild-type, EP4^−/−^, Vav^Cre^EP4^fl/fl^, and their appropriate control mice were anaesthetised by inhalation of isoflurane and the pro-allergic cytokine IL-33 (200 ng per treatment) was administered via intratracheal administration once daily for consecutive 3 d. When indicated, EP2 or EP4 agonist (each 10 μg) or vehicle control (PBS) were co-administered with IL-33. Indomethacin (5 mg/kg/day) or vehicle (0.5% EtOH) was administered in drinking water. 2-Deoxy-d-glucose (2-DG, 1 g/kg/d) (Abcam) was administered via intraperitoneal injection (one injection per day throughout the whole experiment). A.A extract (10 μg) was delivered by intranasal administration together with the EP2 or EP4 agonist (each 10 μg) to wild-type C57BL/6 mice which were then culled 24 h after administration. All animals were culled via anaesthetic overdose by i.p injection of 200 μL Pentoject Pentobarbital Sodium 200 mg/mL solution and subsequent exsanguination. Bronchoalveolar lavage (BAL) was collected (x3 lavages with 800 μL ice cold PBS) followed by aseptic lung dissection for subsequent tissue digestion.

### Murine lung and BAL single cell suspensions

Murine lungs were digested in Liberase TL (Roche) and DNase I (Sigma) at 37°C for 35 min. Digested tissue was passed through a 100 μm cell strainer and red blood cells lysed using ACK lysing buffer. Viable and dead lung cell counts were determined using 0.4% Trypan Blue solution and a TC10™ Automated Cell Counter (BioRad). BAL samples were centrifuged and supernatants from the first BAL retrieval harvested, snap frozen on dry ice and stored at −80°C for subsequent cytokine detection using mouse IL-5/IL-13 ELISA kits according to manufacturer’s instructions. Red blood cells were lysed from BAL cell pellets using ACK lysing buffer and cell counts performed as before.

### Flow cytometry

For surface staining only, lung and BAL single cell suspensions were washed in PBS and stained with LIVE/DEAD™ Fixable Blue Dead Cell Stain Kit. Cells were then blocked with IgG from rat serum and surfaced stained. For lung and BAL neutrophil/eosinophil detection, cell suspensions were labelled with CD45, CD11b, CD11c, Ly6G and SiglecF. For BAL ILC2 detection, cell suspensions were labelled with a cocktail of lineage negative (Lin-) markers consisting of CD3e, CD19, CD11b, CD11c and NK1.1, as well as CD45, CD4, CD90.2, ICOS and ST2. For intracellular staining, single cell lung suspensions were stimulated with a cocktail of PMA, Ionomycin and Brefeldin A (or Golgiplug) and cultured in complete RPMI for 4 h at 37 °C with 5% CO_2_. After stimulation, lung cells were Live/Dead stained, blocked and surface stained for ILC2s as before. Next, lung cells were fixed with intracellular fixation buffer (eBioscience), and then stained for intracellular IL-5, IL-13 and Ki-67 for 20 min at 4 °C. Cell samples were aquired on a BD LSR Fortessa analyser, and results were analysed by FlowJo™.

### ILC2 *in vitro* culture

For *in vitro* experiments all work was carried out aseptically and single cell suspensions were obtained from lungs and bone marrow of untreated *Rag2*^−/−^ mice. Whole lung and bone marrow cells were cultured in complete RPMI with a cytokine cocktail (IL-2, IL-7, IL-33) with various reagents indicated in the figure legends at 37 °C, 5% CO_2_ for 3-4 days. IL-5 and IL-13 in the supernatants were analysed by ELISA. In some experiments, CD90.2^+^ cells were pre-sorted from *Rag2*^−/−^ lung and bone marrow single cell suspensions using an EasySep™ Mouse CD90.2 Positive Selection Kit II (Stemcell Technologies) according to manufacturer’s instructions. Live Lineage(CD11c/CD11b/NK1.1)^−^CD45^+^CD90.2^+^ST2^+^ ILC2s were sorted using a BD FACS Aria II and then cultured in complete RPMI containing 50 μM β-mercaptoethanol with IL-2, IL-7, and IL-33 with or without PGE_2_ in a round bottom 96-well plate at 37 °C in 5% CO_2_ for 3 days.

### Seahorse assays

The Seahorse XFe96-well metabolic analyser (Agilent) was used to investigate the effect PGE_2_ may have upon ILC2 glycolysis *in vitro*. *RAG2^−/−^* mice were subjected to i.t administrations of IL-33 (one i.t. per mouse per day) on days 0, 2 and 4. Lung and BAL single cell suspensions were prepared on day 6 and pooled together for sorting ILC2s as described above. 4 × 10^4^ ILC2s were rested overnight in a 96-well round bottom plate in complete RPMI and IL-2 and IL-7 (both 10 ng/mL). On the next day, IL-33 (10ng/mL), PGE_2_ (1 μM) or DMSO (0.01% v/v as vehicle) was added to media and cultured for a further 24 h. The next day, ILC2 counts were performed on each treatment to confirm >95% viability. Approximately 4 × 10^4^ ILC2s re-suspended in 50 μL Seahorse XF RPMI pH 7.4 (Agilent) (+ 1M glucose, 100 mM pyruvate and 200 mM glutamine) were added in triplicate to wells of a Seahorse XF96 V3 PS cell culture microplate (Agilent) coated 24 h prior with Cell-Tak™ cell adhesive (Sigma). Precise measurements of glycolysis in ILC2s were carried out using the Seahorse XF Glycolytic Rate Assay Kit (Agilent; 103344-100) according to manufacturer’s instructions. The real time oxygen consumption rate (OCR), extracellular acidification rate (ECAR) and proton efflux rate (PER) of ILC2s during glycolysis was directly measured, including glycolytic rates for basal conditions and compensatory glycolysis after injections of 0.5 μM Rotenone/Antimycin A (inhibits mitochondrial respiration) and 50 mM 2-DG (inhibits glycolysis) respectively into all treatment wells.

### RNA-sequencing data analysis

To explore the effect of cAMP on ILC2 gene expression, we first downloaded raw sequence data in FASTQ format from the GEO database (accession number: GSE131996). FastQC was used to assess the quality of FASTQ files (https://www.bioinformatics.babraham.ac.uk/projects/fastqc/). Sequence reads were mapped to mouse genome using STAR 2.7 (54). QualiMap (55) was used to assess the quality of mapped data and featureCounts was employed to count uniquely mapped fragments against genomic features defined by the GENCODE annotation file (Mus_musculus.GRCm39.103.gtf). The counts were further processed for differential expression gene analysis in R (4.0.3). Differentially expressed genes were analysed by DESeq2 (56) and significance was identified using adjusted p value < 0.05 and the absolute value log2 fold change ≥ 1. Ensembl version 103 was used for gene annotation. Gene Set Enrichment Analysis (57) was performed for functional analysis using the Molecular Signatures Database v7.4. Hallmark, KEGG and REACTOME curated gene sets were used as references for screening enriched pathways. *P* values were adjusted and enrichment scores were normalised. An adjusted *P* value < 0.05 (Benjamini–Hochberg) was used as an indicator of significance in pathway analysis.

### Statistical analyses

All data were expressed as mean ± SD (*in vivo*) or SEM (*in vitro*). For certain experiments where both sexes of animals were used, absolute cell numbers were normalised by sex and data were presented as fold changes. Statistical significance between two groups was examined using an unpaired, 2-tailed Student’s *t*-test. One-way and two-way ANOVA with post-hoc Holm-Sidak’s multiple comparisons tests were used to evaluate statistical significance between multiple groups. Statistical analyses were performed using Prism 8 software (Graphpad) and significance was accepted at *P*<0.05.

### Study approval

All animal experiments were conducted in accordance with the U.K. Animals (Scientific Procedures) Act of 1986 with local institutional ethical approval by the University of Edinburgh Animal Welfare and Ethical Review Body.

## Author contributions

CTR and CY conceived of all the experiments. CTR, JMF, MG, and PJ performed the experimental work. YZ and BZ analysed RNAseq data. DJS, RMB, SN, HJM, RMM, JKJS, and AGR provided essential animal lines, key reagents and critical input. The manuscript was written by CTR and CY, with critical input from YZ, RMB, SN, HJM, RMM, JKJS, and AGR. This project was managed and supervised by CY.

## Acknowledgements

We thank E. Dzierzak for providing Vav-Cre mice; P.J. Brophy and D. Mahad for CD90.2(Thy1.2)-iCre mice; and C. Ffrench-Constant for Rag2^−/–^ mice. We also thank F. Rossi, S. Johnston, W. Ramsay, and M. Pattison at the University of Edinburgh QMRI and SCRM flow cytometry facilities for cell sorting and analysis; staff at the University of Edinburgh LFR and SCRM animal facilities for technical support; and C. Elder and M. McIlorum for technical assistance. This work is supported in part by UKRI Medical Research Council (MR/R008167/1 to C.Y.), Cancer Research UK (C63480/A25246 to C.Y.). M.G. received a CMVM Research Adaptation Funding support.

## Supplementary Materials

**Supplementary Figure 1.**
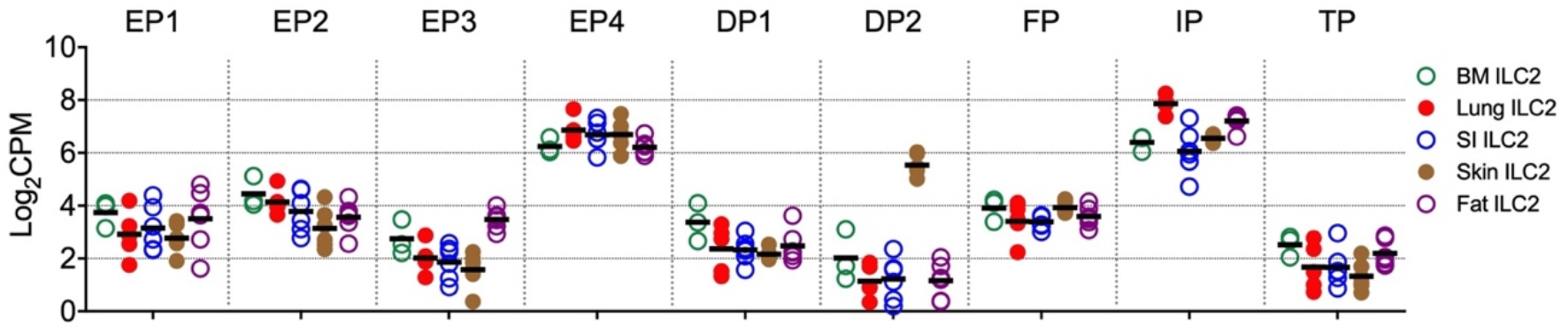
Gene expression of prostaglandin receptors on ILC2s isolated from various tissues. Raw bulk RNAseq data was retrieved from Gene Expression Omnibus GSE117470.

**Supplementary Figure 2.**
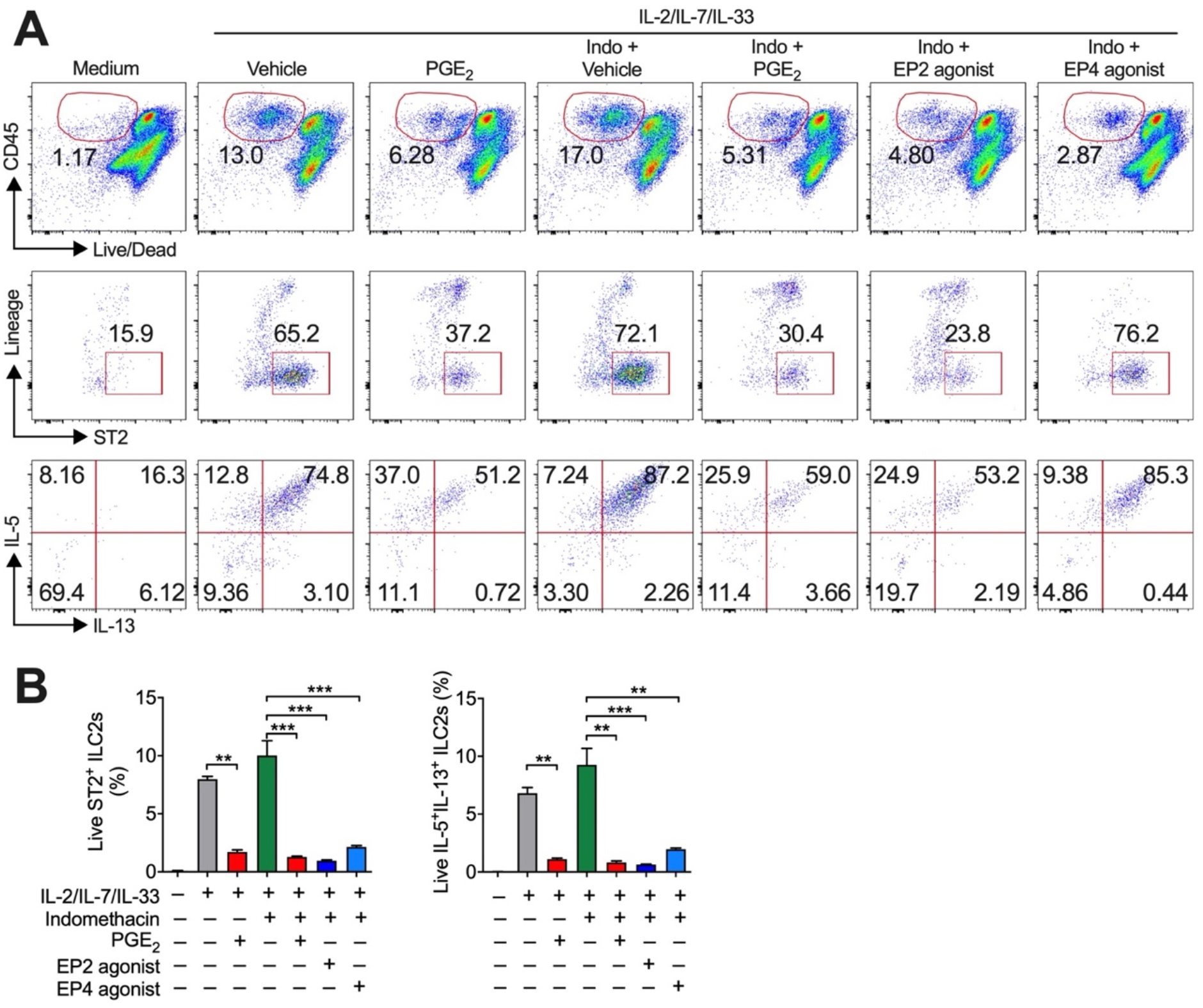
Effects of PGE2 on lung ILC2 responses *in vitro*. Whole populations of single cells isolated from Rag2^−/−^ lungs were cultured with IL-33, IL-2 and IL-7 in the presence or absence of indomethacin (Indo), PGE_2_, EP2 agonist or EP4 agonist. (A) Representative flow cytometric dot plots. **(B)** Collective percentages of total live ST2^+^ or IL-5+/IL-13-expressing ILC2s. Data shown as means ± SEMs are one from 2 experiments. \*\**P*<0.01, \*\*\**P*<0.001, \*\*\*\**P*<0.0001 by one-way ANOVA with post-hoc Holm-Sidak’s multiple comparisons tests.

**Supplementary Figure 3.**
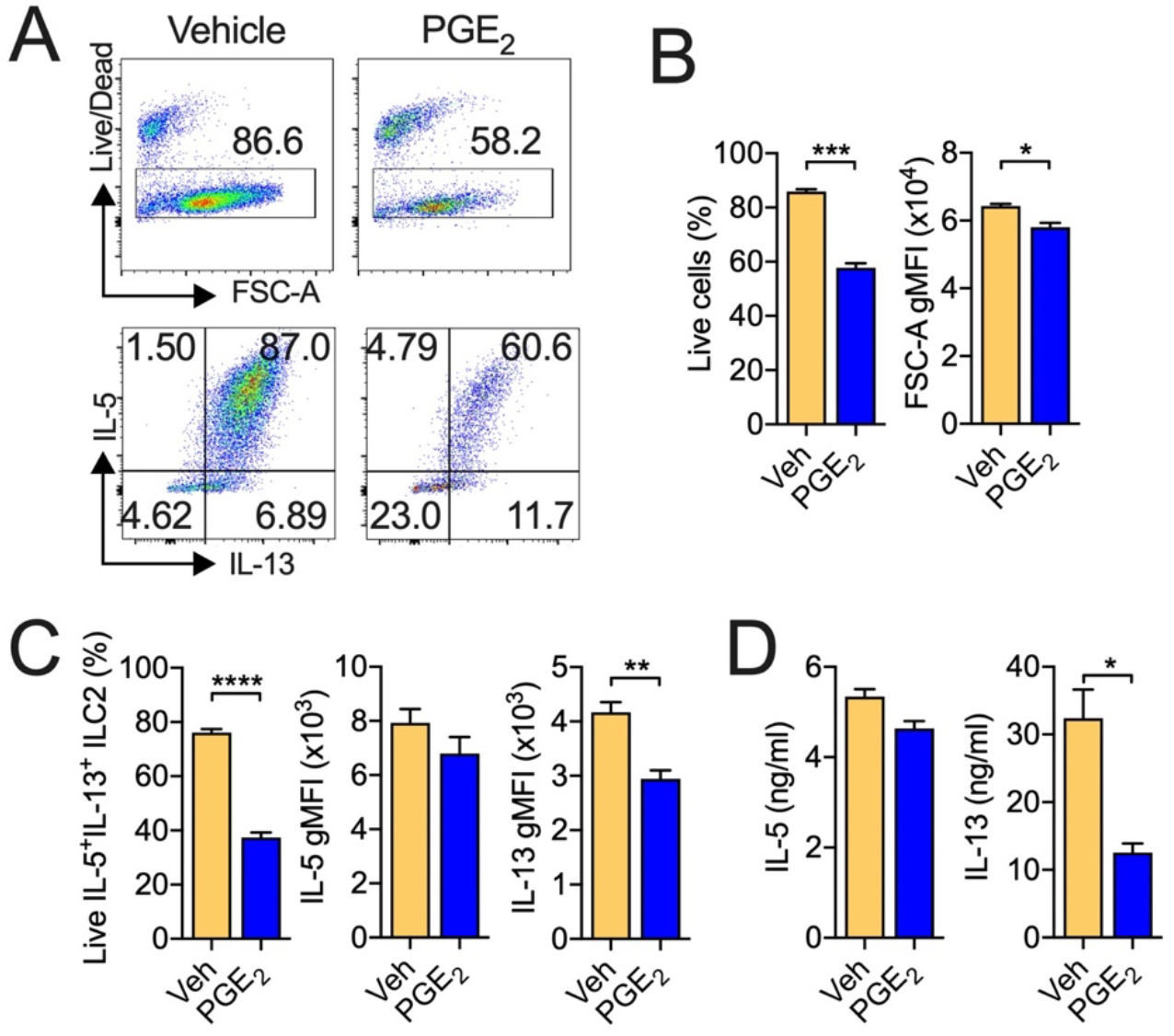
Effects of PGE2 on bone marrow ILC2 responses *in vitro*. ILC2s were sorted from Rag2^−/−^ bone marrow cells and cultured with IL-33, IL-2 and IL-7 in the presence or absence of PGE_2_ for 3 days. **(A)** Representative flow cytometric dot plots. **(B)** Viability and cell size (indicated by FSC-A gMFI). **(C)** Percentages of live IL-5^+^IL-13^+^ ILC2s and gMFI for IL-5 and IL-13. **(D)** Cytokine secretion in the cell culture supernatants. Data shown as means ± SEMs are from one of two independent experiments. \**P*<0.05, \*\**P*<0.01, \*\*\**P*<0.001, \*\*\*\**P*<0.0001 by unpaired, 2-tailed Student *t*-tests. Veh, vehicle.

